# Chromatin accessibility and heat stress gene expression in the reef-building coral, *Acropora millepora*

**DOI:** 10.1101/2025.08.24.672008

**Authors:** Leslie A Guerrero, Kenzie N Pollard, Rachael A Bay

## Abstract

Understanding how organisms regulate gene expression to maintain homeostasis in the face of environmental changes is critical, particularly considering increasing climate stressors. While DNA methylation has dominated the literature on epigenetic control of gene expression in ecological systems, alternative epigenetic layers may provide additional insight into mechanisms of gene expression regulation. This study investigates the role of chromatin accessibility in influencing gene expression in the reef-building coral *Acropora millepora*, especially in response to thermal stress. We find that highly accessible regions of chromatin, or open chromatin, are predominantly located in distal intergenic regions, promoters, and introns. Genes with open promoters exhibit increased expression and reduced variability, suggesting that chromatin accessibility may influence gene expression plasticity. The baseline state of promoter accessibility is weakly correlated with the expression response to heat stress, suggesting chromatin state can influence organismal response to environmental stress. This study contributes new insights into regulatory mechanisms important for responding to acute environmental stressors. This work also establishes a foundation to investigate the interactions between chromatin accessibility, additional epigenetic layers, and how the dynamics of these interacting epigenetic layers contribute to adaptive molecular and cellular responses, which will be critical for understanding organismal resilience to ongoing environmental change.

## Introduction

To maintain homeostasis, organisms must effectively respond to external cues at both the cellular and molecular levels. A critical aspect of this response is the regulation of gene expression, which ultimately contributes to shifts in phenotypes. The mechanisms by which gene expression is precisely mediated to respond to complex environmental signals remain a major biological question. Epigenetics provides a valuable lens for understanding how environmental factors influence gene expression, leading to functional trait variation in natural populations (Kilvitis et al. 2014; Lamka et al. 2022). Defined as the study of molecular mechanisms that interact with DNA to modulate gene expression without altering the underlying sequence, epigenetics explores processes that play a crucial role in how organisms respond to biotic and abiotic factors such as temperature fluctuations and predator cues (Feil and Fraga 2012; Noguera and Velando 2019). For example, in plants, abiotic stressors, such as drought, influence changes in non-coding RNA expression, which is an epigenetic mechanism that regulates coding gene expression in a stress-specific manner (Di et al. 2014). Epigenetic mechanisms are also important for seasonal responses, reproductive trait fluctuations, and generating phenotypic variation in clonal species (Fishman and Tauber 2024; Jueterbock et al. 2020; Lindner et al. 2020).

A core objective of ecological epigenomics is to unravel the intricate interactions between epigenetic layers, gene expression regulation, and their ecological and evolutionary significance (Lamka et al. 2022). This endeavor has become increasingly important for predicting species’ adaptive potential as climate change intensifies environmental stressors, particularly for long-lived species that cannot easily migrate to more favorable habitats (Eirin-Lopez and Putnam 2019; Hofmann 2017; Lancaster et al. 2022). Reef-building corals, which are highly vulnerable to ocean warming and other stressors, present a compelling system for studying the role of epigenetic mechanisms in regulating gene expression in response to abiotic stressors. Corals possess an ability to modulate their gene expression in response to acute environmental stress and, following conditioning, have the capacity to finely-tune their gene expression for impending stressors (Bay and Palumbi 2015; Bellantuono et al. 2012). This suggests that molecular mechanisms can preserve the ‘memory’ of exposure to environmental cues and facilitate the necessary gene expression response to maintain homeostasis.

DNA methylation is the most extensively studied epigenetic mechanism in corals. Across various coral taxa, DNA methylation fluctuates in response to environmental changes (Dimond and Roberts 2016; Dimond and Roberts 2020; Dixon et al. 2018; Putnam et al. 2016; Rodriguez-Casariego et al. 2022). However, accumulating evidence indicates that shifts in DNA methylation following environmental fluctuations may play a negligible role in mediating gene expression changes (Abbott et al. 2024; Dixon and Matz 2022; Guerrero and Bay 2024; Rodriguez-Casariego et al. 2022; but see Gomez-Campo et al. 2023; Rodríguez-Casariego et al. 2020). Despite this, DNA methylation has established roles in suppressing transposable elements, maintaining transcriptional homeostasis, and correlating with higher overall gene expression (Dixon and Matz 2022; Guerrero and Bay 2024; Li et al. 2018; Rodriguez-Casariego et al. 2022; Ying et al. 2022). Altogether, DNA methylation in corals is hypothesized to act as a regulatory mechanism over adaptive timescales rather than a change in response to short-term environmental changes (Abbott et al. 2024). DNA methylation studies have revealed fascinating aspects of coral biology, but other regulatory layers remain unexplored in understanding the coral epigenetic landscape.

Emerging interest in chromatin organization and accessibility has provided new insights into this epigenetic layer as a mediator of the stress response in cnidarian species, including corals (Rodriguez-Casariego et al. 2018; Roquis et al. 2022; Weizman and Levy 2019). Eukaryotic DNA is organized and tightly packed in the nucleus through physical interactions with octamer cores of histone proteins (Tsompana and Buck 2014). Regions of open chromatin result from post-translational histone modifications or histone variant substitutions that alter nucleosome structure, thereby facilitating DNA-binding processes such as transcription activation, DNA repair, and recombination (Tsompana and Buck 2014). Recent studies have shown that coral histones undergo post-translational modifications following stress events. For example, in *Acropora cervicornis*, phosphorylation of the histone variant H2.AX following nutrient and heat stress activates DNA repair processes (Rodriguez-Casariego et al. 2018). Similarly, in *Pocillopora acuta*, acute heat stress leads to clipping of canonical histone H3 amino acid tails, although the functional consequences of this modification remains to be explored (Roquis et al. 2022). Notably, studies in ATAC-seq and RNA-seq in the cnidarian model species *Exaiptasia pallida* have revealed that chromatin accessibility dynamics are influenced by thermal exposure, with exposed transcription factor binding motifs being linked to gene expression changes (Weizman and Levy 2019). However, the relationship between baseline chromatin accessibility and gene expression in reef-building corals remains unexplored, a crucial step to enhance our understanding of epigenetic regulation in these ecologically vital species.

In this study, we use ATAC-seq to investigate regions of open chromatin and their relationship to gene expression variation broadly and to the heat stress gene expression response in the reef-building coral, *Acropora millepora*. Specifically, we subjected *A. millepora* fragments to an acute heat stress assay and performed 3’TagSeq to evaluate the heat stress gene expression response. In a separate set of fragments, we performed ATAC-seq, which we used to assess the relationship between promoter accessibility, overall expression and expression variation. Finally, we investigated whether the accessibility of chromatin promoters influences gene expression during heat stress. The relationship between open chromatin promoters and gene expression established here, through the integration of ATAC-seq and 3’TagSeq, sets the stage to broaden our understanding of mechanisms driving gene expression changes in corals.

## Methods

### Experimental Design

#### Establishing thermal response

We acquired 74 fragments of *Acropora millepora*, comprised of 8 different genotypes, from Neptune Aquatics, Inc. in San Jose, California. Because we are most interested in identifying a core transcriptomic stress response, we used these fragments to build a library of samples with varying thermal histories in an attempt to establish a general heat stress response. The fragments underwent a 10-day conditioning period in the laboratory aquarium, which was maintained at 24.5°C. We maintained the salinity at 32 ppm as measured with a refractometer. To vary thermal exposure history, we incubated 24 fragments at 27°C for 7 days while keeping another 24 fragments at the control temperature of 24.5°C, then returned all fragments to 24.5°C. Due to space limitations, we could not include fragments from all genotypes in every acclimation treatment. However, we ensured each genotype was represented in its respective thermal acclimation and control treatments. To characterize the thermal response, we performed five heat stress assays over the course of three weeks using a modified Coral Bleaching Automated Stress System (CBASS) protocol (Voolstra et al. 2020). The heat stress treatment involved a 3-hour thermal ramp from 24.5°C to 36°C, followed by a 3-hour hold at 36°C to simulate thermal stress conditions, and finally, thermal decline over the course of an hour back to 24.5°C. Temperature profiles were picked based on pilot experiments that tested for a temperature that induced an intermediate bleaching response that could capture variation among coral fragments but did not result in tissue death. Heat stress assays were performed on 5 genotypes on Day 0 to capture baseline thermal tolerance (Figure 1). We followed with assays on days 7, 11, 14, and 21 on 4 fragments from 4 different genotypes with a thermal acclimation treatment and their respective controls (Figure 1). At the end of the heat stress assay, we dark-acclimated the fragments for an hour and measured the efficiency of photosystem II (Fv/Fm) of each fragment using Pulse Amplitude Modulated (PAM) fluorometry. We used a Wilcoxon rank-sum test to compare the distributions of Fv/Fm values between control and heated samples.

**Figure 1.**
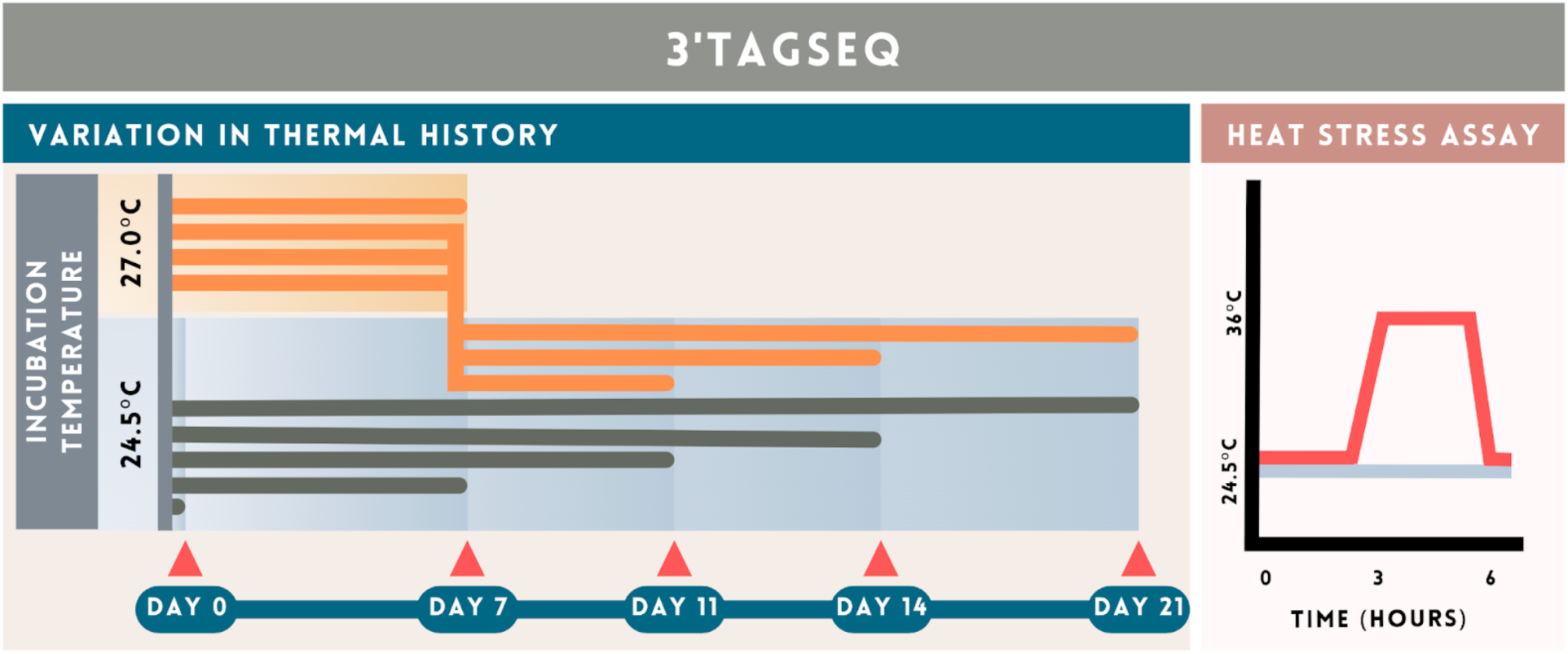
Experimental design. The left panel illustrates treatments designed to induce variation in thermal history. Fragments held at 24.5°C are the thermal history controls. Triangles indicate time points for heat stress assay. The right panel depicts heat stress assay temperature profile. Heat stress assays were conducted on fragments with varying thermal histories, alongside thermal history controls. We preserved tissue from this experiment for 3’TagSeq only (i.e., ATAC-seq was not performed on any of these fragments).

After the PAM fluorometry, we captured images of the fragments with Kodak color standards using an iPhone (Kodak color separation guide and grayscale Q-13, Kodak, USA). We measured the red channel intensity using the methods described by Winters et al. (2009). We used a t-test to compare the average red channel intensity values between control and heated samples. Within an hour of completing PAM fluorometry measurements and color-standard photography, we flash-froze the fragments and stored tissue samples at -80°C until RNA and protein extraction.

We assayed the activity of glutathione reductase (GR), an emerging indicator of coral stress, for control and heat stress samples (Majerová and Drury 2022). In brief, we homogenized coral tissue by vortexing 1g of thawed coral clipping with 0.1mL of glass beads (Sigma-Aldrich, Mfr. Catalog ID g8772-10g) and 1 mL of filtered seawater (sl35) in a microcentrifuge tube. We used compound microscopy to check that symbiont cell integrity was maintained, thus minimizing the risk of symbiont GR contamination. We isolated coral tissue from symbionts by centrifuging samples at 800 g for 5 minutes and removed cellular debris by centrifuging the supernatant containing coral cells at 14,000g for 10 minutes. We collected the supernatant and quantified total protein using the Broad Range Qubit assay (Thermo Fisher Scientific, Mfr Catalog ID A50668). We used the EnzyChrom Glutathione Reductase kit (BioAssays, Mfr Catalog ID ECGR-100) to measure GR absorbance and calculate GR activity following manufacturer instructions. We used a Spectramax M2 plate reader (Molecular Devices, San Jose, CA) to measure absorbance at 412 nm. We normalized the GR activity with the total protein concentration for each sample. We used a Wilcoxon rank-sum test to compare the distributions of normalized GR activity values between control and heat-stressed samples.

### Molecular Preparation and Sequencing

#### 3’TagSeq

We used 3’TagSeq to profile gene expression responses to heat stress. Our samples, while all exposed to similar heat stress, comprised a range of thermal histories. We extracted RNA from 58 samples using the E.Z.N.A. HP Total RNA Isolation Kit (Omega Bio-Tek, Mfr. Catalog ID R6812) using instructions and the DNase I Digestion Protocol supplied by the manufacturer. We stored extracted RNA at -80°C. We sent RNA to the DNA Technologies and Expression Analysis Core at the UC Davis Genome Center for library preparation and sequencing. Barcoded 3’Tag-Seq libraries prepared using the QuantSeq FWD kit (Lexogen, Vienna, Austria) for multiplexed sequencing according to the manufacturer’s recommendations. The fragment size distribution of the libraries was verified via micro-capillary gel electrophoresis on a Bioanalyzer 2100 (Agilent, Santa Clara, CA). The libraries were quantified by fluorometry on a Qubit instrument (LifeTechnologies, Carlsbad, CA), and pooled in equimolar ratios. Fifty-eight libraries were sequenced on one lane of an AVITI sequencer (Element Biosciences, San Diego, CA) with single-end 100 bp reads. The sequencing generated more than 4 million reads per library.

We trimmed the reads of the universal Illumina adapters and bases on the 3’ end of the read with a sequencing quality score of less than 10 using Cutadapt (v. 4.1) (Martin 2011). To separate intracellular symbiont reads from coral reads, we used BBsplit (v. 38.22) to bin reads that align to either the *Symbiodinium goreaui* or the *A. millepora* genome (reefgenomics.org; Bushnell 2014; Fuller et al. 2020). To optimize TagSeq read binning, we adjusted the BBsplit maxindel argument to ‘100K’. We mapped the *A. millepora* reads to the *A. millepora* reference genome using the STAR aligner (v. 2.7.10b) (Dobin et al. 2013). We used HTSeq (v. 2.0.3) to count all the reads that mapped to genes, including reads that mapped to multiple locations (--nonunique all) using the annotated gene models provided with the *A. millepora* reference genome (Anders et al. 2015; Fuller et al. 2020).

#### ATAC-seq

We assayed ‘open,’ or hyper-accessible, chromatin in *A. millepora* using ATAC-seq (Buenrostro et al. 2015). Our initial plan was to conduct ATAC-Seq on the same fragments used for gene expression analysis. However, due to failed reactions, we were unable to use these same samples. Instead, our study explores a more general connection between chromatin accessibility and gene expression in *A. millepora.* We acquired four fragments of *A. millepora*, each of a different genotype, from Neptune Aquatics, Inc. in San Jose, California. The fragments were stored in the laboratory aquarium overnight at 24.5°C. The following morning, we isolated nuclei and prepared DNA for sequencing regions of open chromatin by modifying previously published protocols (Ackermann et al. 2016; Buenrostro et al. 2015; Weizman and Levy 2019). The fragments used for ATAC-seq were not subject to any heat stress as we focused on open chromatin regions at baseline conditions. We first isolated tissue from the skeleton by scraping the tissue with a sterile scalpel into ice-cold 1x PBS buffer. We placed a sterile Petri dish on ice for each sample, then slowly pipetted the tissue and PBS buffer mixture up and down 10 times in the dish to evenly distribute the tissue. We pipetted 750 µL of the mixture into a 40 µm mini cell strainer fit into a 1.5 mL microcentrifuge tube to isolate coral cells. After collecting the filtrate, we centrifuged the sample tubes at 1500g for 5 minutes at 4°C. We pipetted off the supernatant and resuspended the isolated cells in 500 µL of fresh ice-cold 1x PBS buffer. To pellet cells with symbionts, we centrifuged the samples at 600g for 10 minutes at 4°C. We collected and resuspended the supernatant in fresh ice-cold 1x PBS. We centrifuged the sample tubes at 1500g for 5 minutes at 4°C and pipetted off the supernatant.

To isolate the nuclei from the total cellular debris, we resuspended the cell pellet in 50 µL of lysis buffer consisting of 0.25 µL of 10% NP40 added to 49.75 µL of resuspension buffer (10 mM Tris-HCl, 10 mM NaCl, 3 mM MgCl_2_). We incubated the cells in the lysis buffer on ice for 3 minutes. Following the 3-minute incubation, we centrifuged the mixture at 300g for 10 minutes at 4°C. We removed the supernatant with cellular debris and repeated the 3-minute incubation in the lysis buffer followed by the 10-minute centrifuge. We removed the supernatant and suspended nuclei in 100 µL of fresh ice-cold resuspension buffer. We centrifuged this mixture at 300g for 10 minutes at 4°C. We removed the supernatant and resuspended the nuclei in 100 µL fresh ice-cold resuspension buffer. To estimate the number of nuclei we collected per genotype, we stained 10 µL of suspended nuclei with DAPI to visualize and count nuclei using a hemocytometer and fluorescent microscopy (Life Technologies EVOS FL Color, Mfr Catalog ID AMEFC4300).

To perform the tagmentation, we added 2.5 µL of Tn5 Transposase suspended in 1X TD Buffer (Nextera DNA sample preparation kit; Illumina, cat. no. FC-121-1030), for every 50,000 nuclei. We incubated this reaction for 30 minutes at 37°C. We purified the DNA using the MinElute Reaction Cleanup Kit (Qiagen, Mfr. Catalog ID 28204).

We performed 20 cycles of PCR using the NEBNext High-Fidelity 2X PCR Master Mix (NEB, Mfr. Catalog ID M0541S) to generate sequencing libraries. We purified the libraries using AMPure XP beads (Beckman Coulter, Mfr. Catalog ID A63880). We used the Agilent High Sensitivity DNA Bioanalysis Kit (Agilent, Mfr. Catalog ID 5067-4626) to assess the quality of the purified libraries and that the expected fragment sizes of amplified ATAC-seq libraries were present (Buenrostro et al. 2015). We pooled the purified libraries for sequencing at the DNA Technologies and Expression Analysis Core at the UC Davis Genome Center. DNA was sequenced using Illumina NextSeq to generate 75 bp PE reads intending to achieve 120 million read pairs total.

We used Cutadapt (v. 4.1) to trim Nextera adapter sequences and bases with a sequencing quality score less than 20 from the 3’ end of the reads (-q 20) (Martin 2011). We used the mitochondrial genome supplied with the *A. millepora* reference genome to distinguish and exclude mitochondrial from nuclear reads using BBsplit (v. 38.22) (Bushnell 2014; Fuller et al. 2020). We mapped reads to the *A. millepora* reference using Bowtie2 (v. 2.4.4) (Langmead and Salzberg 2012). We used samtools view (v. 1.13.0) to filter out reads that were not the primary alignment, PCR or optical duplicates, reads that failed platform/vendor quality checks, and supplementary alignments (-F 3840) (Li et al. 2009). We removed duplicate reads with Picard MarkDuplicates (v. 2.23.2).

We called hyper accessible chromatin, or peaks, with MACS2 (v. 2.2.6) which reports the Poisson distribution p-values used to identify peaks (Zhang et al. 2008). To generate a unified set of open chromatin regions across the four replicates, we used the DiffBind R package (v3.12.0; Stark and Brown 2012). A differential binding affinity (DBA) object was created from a sample metadata sheet, and a consensus peak set was defined using the dba.peakset() function, requiring that peaks be present in at least two out of the four *A. millepora* genotypes (minOverlap = 2). To ensure inclusion of high-confidence peaks, we applied a -log10(p-value) filter of 4.3, corresponding to a peak significance threshold of 5 × 10⁻⁵. We used CHIPseeker (v1.38.0) and the *A. millepora* gene annotation file derived from the *A. millepora* genome assembly v2.01 to annotate the consensus peak set (n=11,231) (Fuller et al. 2020; Wang et al. 2022). To analyze the functional enrichment of open chromatin promoters, we used Gene Ontology (GO) terms with the R package, topGO v. 2.50.0, comparing peak enrichment to all annotated genes in *A. millepora* (Alexa and Rahnenfuhrer 2022, Fuller et al. 2020).

### Heat stress treatment effect on gene expression

We used the R Package DESeq2 v. 1.36.0 (Love et al. 2014; R Core Team 2023) to test the effects of thermal history, genotype, and heat stress on gene expression in our samples. To visualize these effects, we first performed principal component analysis (PCA) and estimated the variation explained by covariates using the R package variancePartition v. 1.32.5 (Hoffman and Schadt 2016). Next, we identified the genes with a significant change in expression due to heat stress. First, we created a ‘DESeqDataSet’ object, filtered for genes with a depth of at least 10 reads in all samples, and normalized data with the variance stabilizing transformation. We defined “heat stress genes” with an absolute value of the log_2_ fold change greater than 2 and with an adjusted p-value less than 0.05. To analyze the functional enrichment of heat response genes, we analyzed Gene Ontology (GO) terms with the R package, topGO v. 2.50.0, comparing the set of heat response genes against a background of all genes that passed our quality filters. Enriched GO terms were identified with the classic Fisher’s test with a p-value of less than 0.01 and at least 10 transcripts within each category (Alexa and Rahnenfuhrer 2022, Fuller et al. 2020).

### Testing the relationship between open chromatin promoters and gene expression

To investigate how chromatin accessibility influences gene expression, we integrated ATAC-seq and RNA-seq datasets. We linked consensus peaks to genes using the gene annotations described above and calculated promoter accessibility by averaging CPM-normalized read counts across the four samples. These counts were generated from the DiffBind object using the edgeR v. 4.0.16 package. Accessibility values were log10-transformed to stabilize variance for visualization and statistical comparison.

We examined whether promoter accessibility varied as a function of gene expression level. To do this, we assigned genes to quintiles based on their DESeq2-calculated baseMean expression values, with the fifth quintile representing the highest expression. Genes with promoter-proximal open chromatin were selected using ChIPseeker annotation (“promoter”), and accessibility profiles were plotted as a function of distance to the transcription start site (TSS) using geom_smooth() in ggplot2 v3.5.1

To assess whether promoter accessibility was associated with gene expression variability, we computed the standard deviation of DESeq2-normalized expression counts across all RNA-seq samples. One outlier gene with extreme variance (standard deviation > 20,000) was removed. We then calculated the coefficient of variation to account for expression magnitude and divided genes into quintiles of expression variability. Genes in the first quintile were categorized as “stable,” while those in the fifth were considered “variable.” We then compared promoter accessibility profiles for stable vs. variable genes, again using smoothed plots of accessibility versus distance to the TSS.

Our primary hypothesis was that gene promoters are poised to facilitate an imminent transcriptional response to heat stress, presumably by histone modifications that yield open chromatin and regulatory protein binding. Using the previously defined set of heat-responsive genes and promoter accessibility data, we tested for enrichment of open chromatin using a Fisher’s exact test. To further examine the relationship between chromatin accessibility and the expression response, we assessed the association between log-transformed promoter accessibility and both mean expression and the absolute value of log2 fold change using Spearman correlation.

## Results

### Phenotypic responses to heat stress

Physiological measures show the expected disruption to symbiosis in response to heat stress. We found a significant decrease in algal symbiont photosystem II efficiency in heat-stressed fragments (Figure 2A, Wilcoxon test, p ≤ 0.0001). Additionally, heat-stressed fragments had a higher visual bleaching response, indicated by a higher red channel intensity–a measure of chlorophyll pigment degradation (Figure 2B, t-test, p ≤ 0.0001). Across all heat stress assays, the heat-stressed fragments have significantly reduced glutathione reductase activity compared to the controls(Figure 2C, Wilcoxon test, p ≤ 0.01). Glutathione reductase activity, an emerging biomarker for coral health, reduces reactive oxygen species (ROS) to a peroxide byproduct, thereby mitigating the downstream deleterious effects on DNA homeostasis caused by ROS (Majerová and Drury 2022). These results collectively indicate that heat stress is detrimental for both the symbionts and coral’s homeostasis.

**Figure 2.**
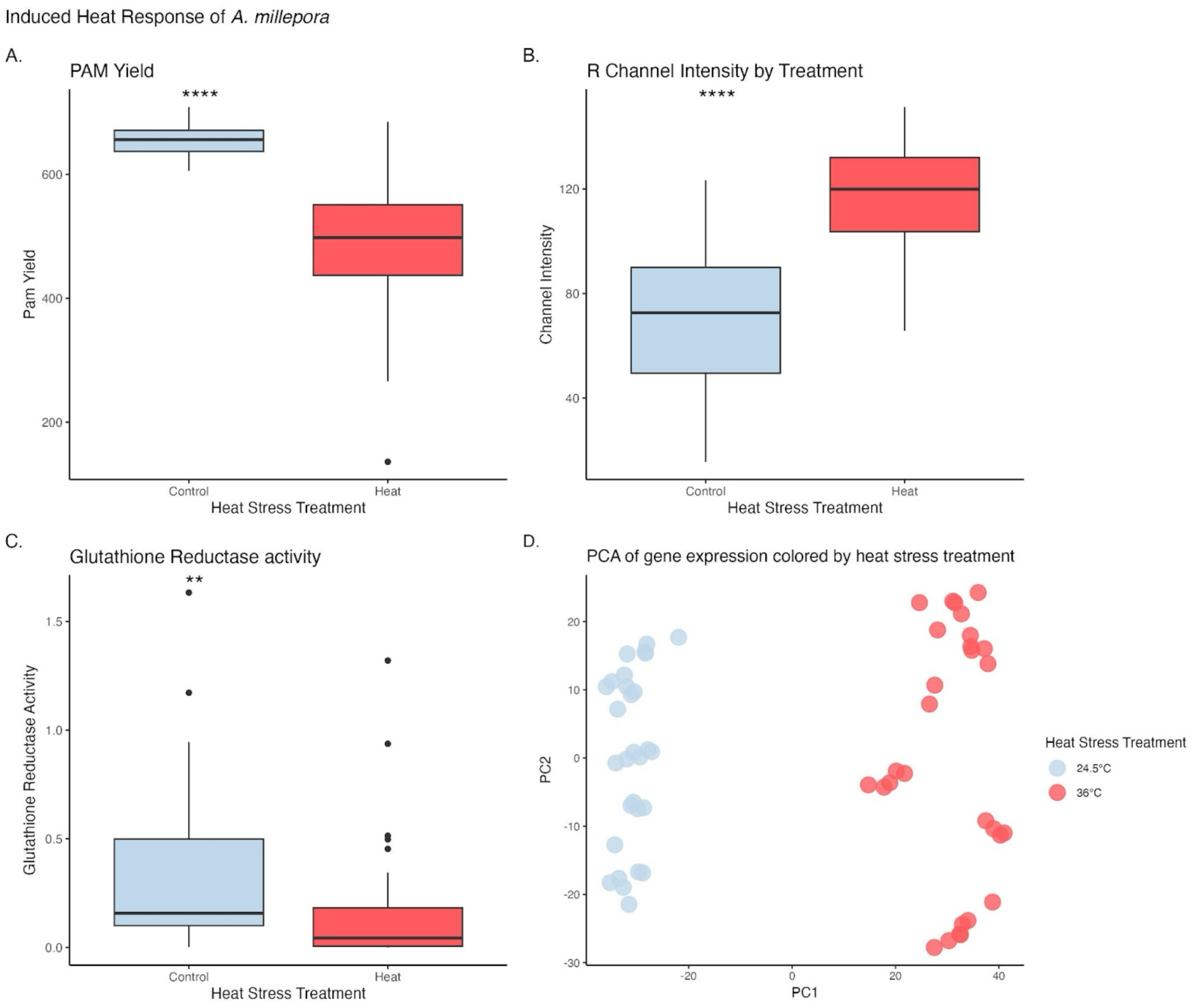
**A. Boxplot of Fv/Fm Yield** showing a reduction in Fv/Fm (Photosystem II yield of algal symbiont) in heat-stressed *A. millepora* fragments compared to control fragments. **B. Boxplot of R channel intensity** showing an increased R channel intensity (higher visible bleaching score) in heat-stressed *A. millepora* fragments compared to the control fragments. **C. Boxplot of glutathione reductase (GR) activity** showing reduced GR activity in heat-stressed *A. millepora* fragments have reduced GR activity compared to the control fragments. **D. PCA of gene expression** in heat-stressed and control fragments of *A. millepora* reveals distinct clustering along the PC1 axis, separating the heat-stressed fragments from the controls.

### Transcriptional response to heat stress

We used RNA 3’TagSeq to evaluate the transcriptomic response to heat stress in *A. millepora*. Reads were binned based on alignment to either the *Symbiodinium goreaui* or the *A. millepora* genome. On average, 1,201,288 reads were derived from symbiont transcripts, and an average of 2,327,833 reads were from *A. millepora* transcripts. We observed an average of 61.3% uniquely mapped coral-derived reads to the *A. millepora* reference genome.

We used DESeq to assess the transcriptional response to heat stress. After filtering for genes with at least 10 reads across all samples, we evaluated the transcriptional response for 19,649 genes. We found that heat treatment is the largest source of variation, followed by genotype (Figure 2D, Supplement Figure 1). Thermal history explained negligible variation in the gene expression response to heat (Supplement Figure 1). Comparing heat stressed to control fragments, 2,227 genes were differentially expressed. Among these heat response genes, 268 GO terms were significantly enriched, with cellular pathways for oxidative stress, unfolding protein binding, transcription, immunity, and apoptosis being particularly notable (Supplement Table 1). These GO terms align with previous reports of *A. millepora* heat stress, where genes associated with oxidative stress, symbiont detection, symbiosis maintenance, and transcription are differentially expressed (Bellantuono et al. 2012; Granados-Cifuentes et al. 2013). These processes are also well documented to be associated with heat stress in other reef-building coral species (Louis et al. 2017).

### ATAC-seq

We successfully performed ATAC-seq on *A. millepora.* After trimming raw reads, we obtained an average of 42,586,471 paired-end reads across all four replicates. On average, 4.8% of reads were derived from mitochondria, which is a low and acceptable amount of mitochondrial contamination (Montefiori et al. 2017). An average of 66.3% of reads were derived from nuclear DNA, and we achieved a primary read alignment rate of 90.1%.

Using MACS2 for peak calling, we identified an average of 32,206 peaks across the four genotypes, with 11,231 consensus peaks identified as overlapping regions of accessibility detected in at least two of the four *A. millepora* genotypes. This number of consensus peaks is lower than the open chromatin regions reported in other invertebrate ATAC-seq studies, possibly reflecting the conservative nature of our peak summary approach (Weizman and Levy 2019; Gatzmann et al. 2018). We found that 50.05% of the open chromatin regions were distal to genic regions (Figure 3). Genic open chromatin occurred more frequently in introns relative to exons (Figure 3), consistent with what is seen in *Strongylocentrotus purpuratus* (Bogan et al. 2023). The relatively high representation of introns among genic peaks underscores the growing recognition of their role in regulating gene expression (Rose 2018; Bogan et al. 2023).

**Figure 3.**
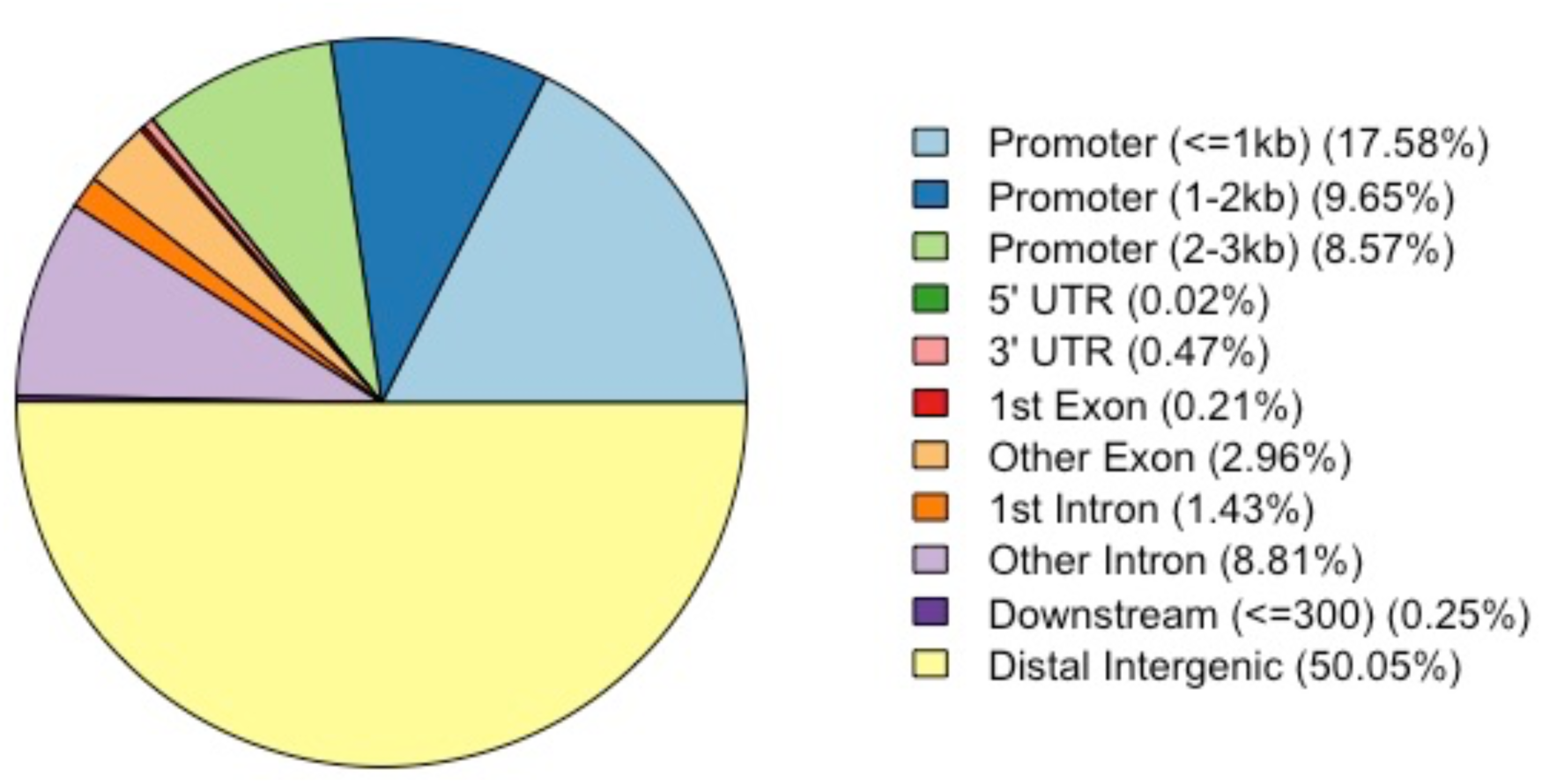
Annotated open chromatin regions in *A. millepora*. Distal intergenic regions compose 50.05% of the open regions, and promoters compose 35.80% of the open regions.

Accessible gene promoters interact with gene regulatory proteins that control transcription initiation and regulation (Danino et al. 2015; Yang and Hansen 2024). Approximately 36% percent (n= 3973) of the consensus peaks were annotated as promoters, which is higher than the typical 25% observed in other studies (Yan et al. 2020). This might indicate that the regulatory regions in our samples were more active. Open chromatin promoter GO terms were linked to critical biological processes that regulate a wide range of molecular functions and cellular activities (Table 1). Specifically, these terms are predominantly associated with immune system regulation, viral interactions and responses, gene expression and transcription regulation, cellular processes and metabolism, and regulation of development and differentiation (Table 1). This enrichment supports the expectation that open chromatin promoters play a key role in higher-level gene expression regulation (Klemm et al. 2019).

**Table 1.**
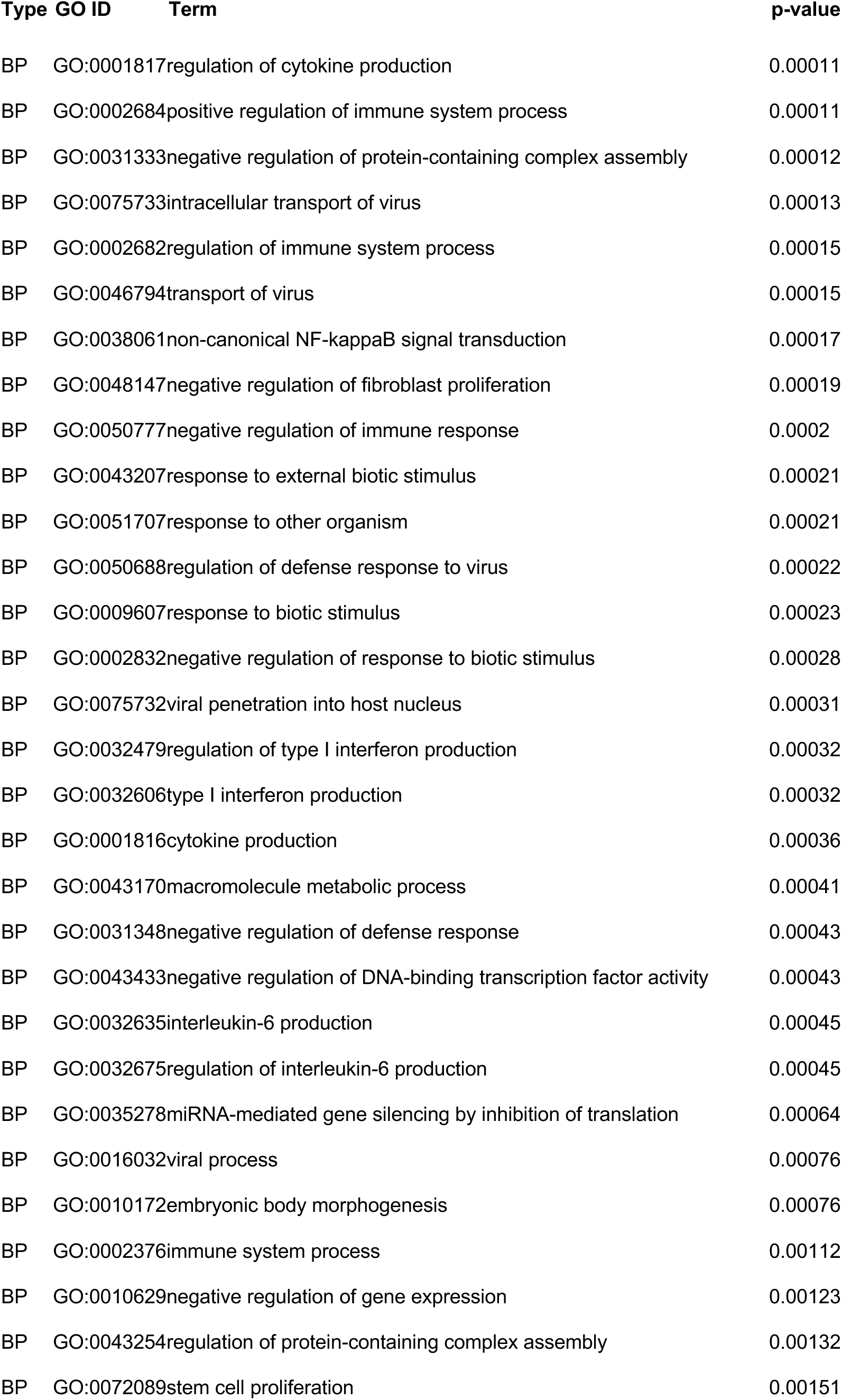
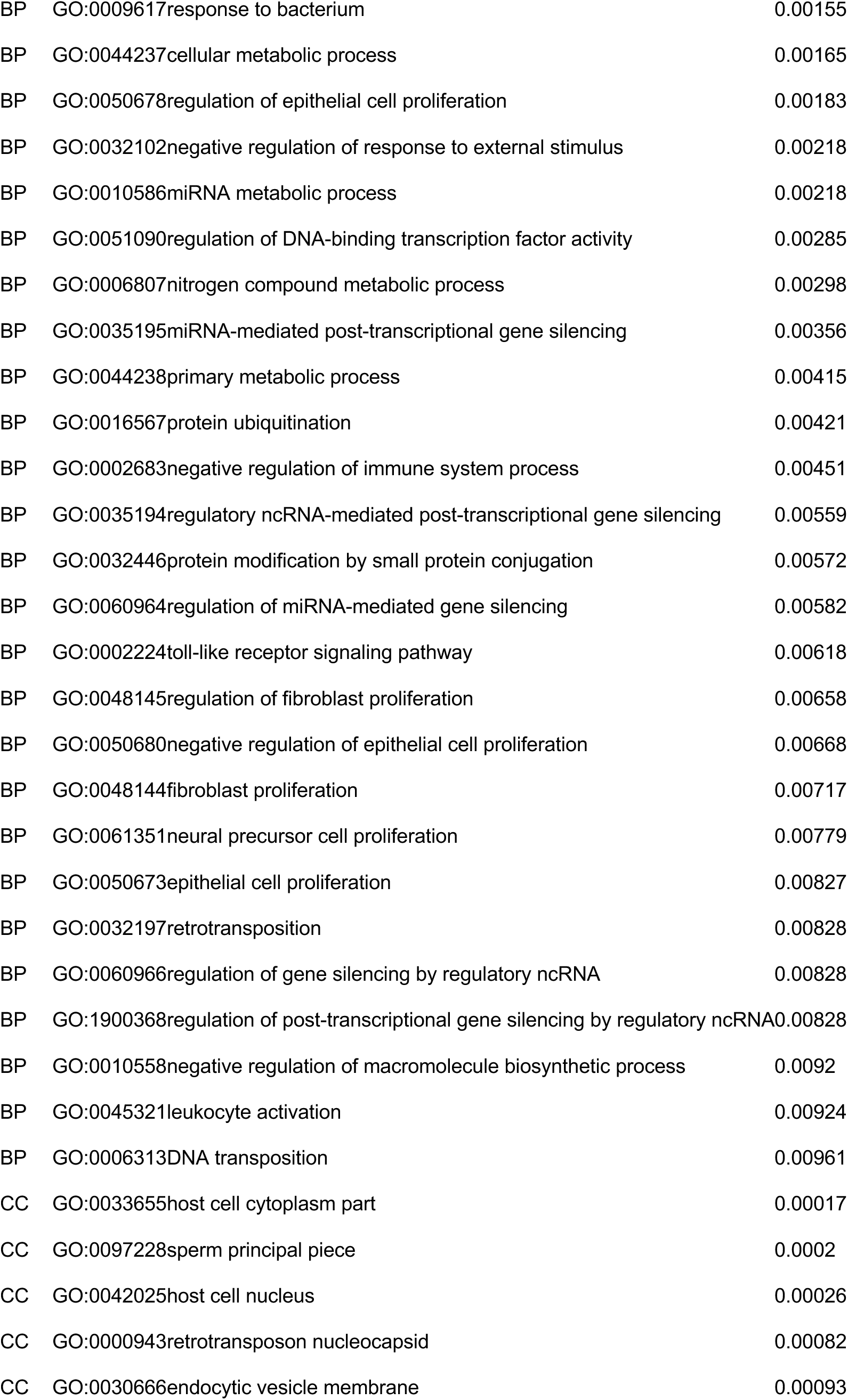
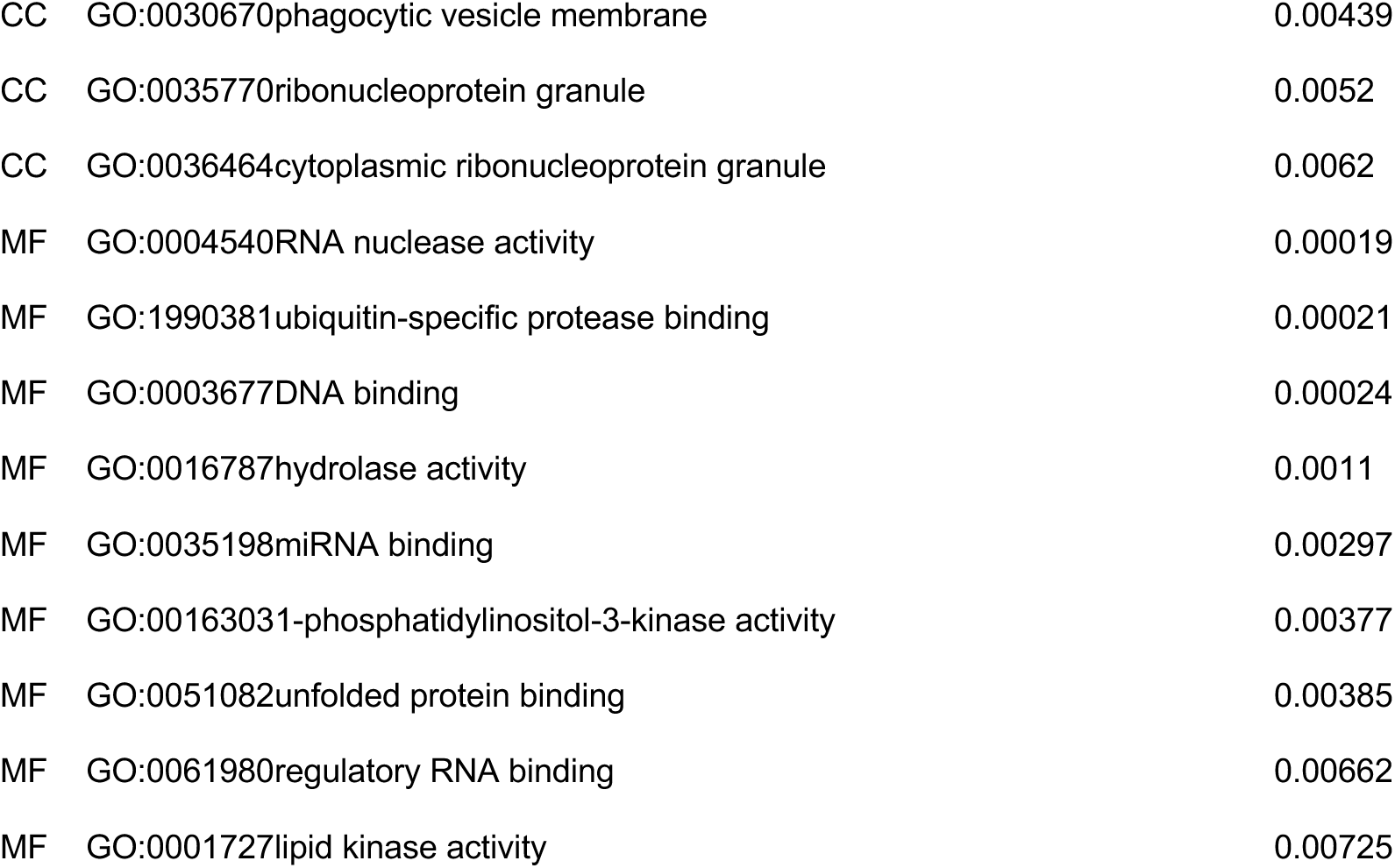
Gene Ontology (GO) terms of open chromatin promoters. Table shows the results from a GO enrichment analysis of open chromatin promoters using the R package, topGO v. 2.50.0. We compared the list of open chromatin promoters in the consensus against a background of all *A. millepora* annotated transcripts. Enriched GO terms were identified with the classic Fisher’s test with a p-value < 0.01 and at least 10 genes within each category. GO terms are contained within ontology types: Biological Processes (BP), Cellular Components (CC), and Molecular Functions (MF).

### Testing the relationship between accessible chromatin promoters and gene expression

We investigated the genome-wide effects of chromatin hyper-accessibility on gene expression. We observed that 1,919 genes in our expression data had open chromatin promoters after collating the consensus peak set with gene expression data. Genes with the highest expression (5th quintile) have more accessible chromatin around the TSS than genes with lower expression (Figure 4A). Genes with more accessible chromatin promoters exhibited the least expression variation (Figure 4B). This is consistent with observations of low-methylated genes in crayfish (Gatzmann et al. 2018).

**Figure 4.**
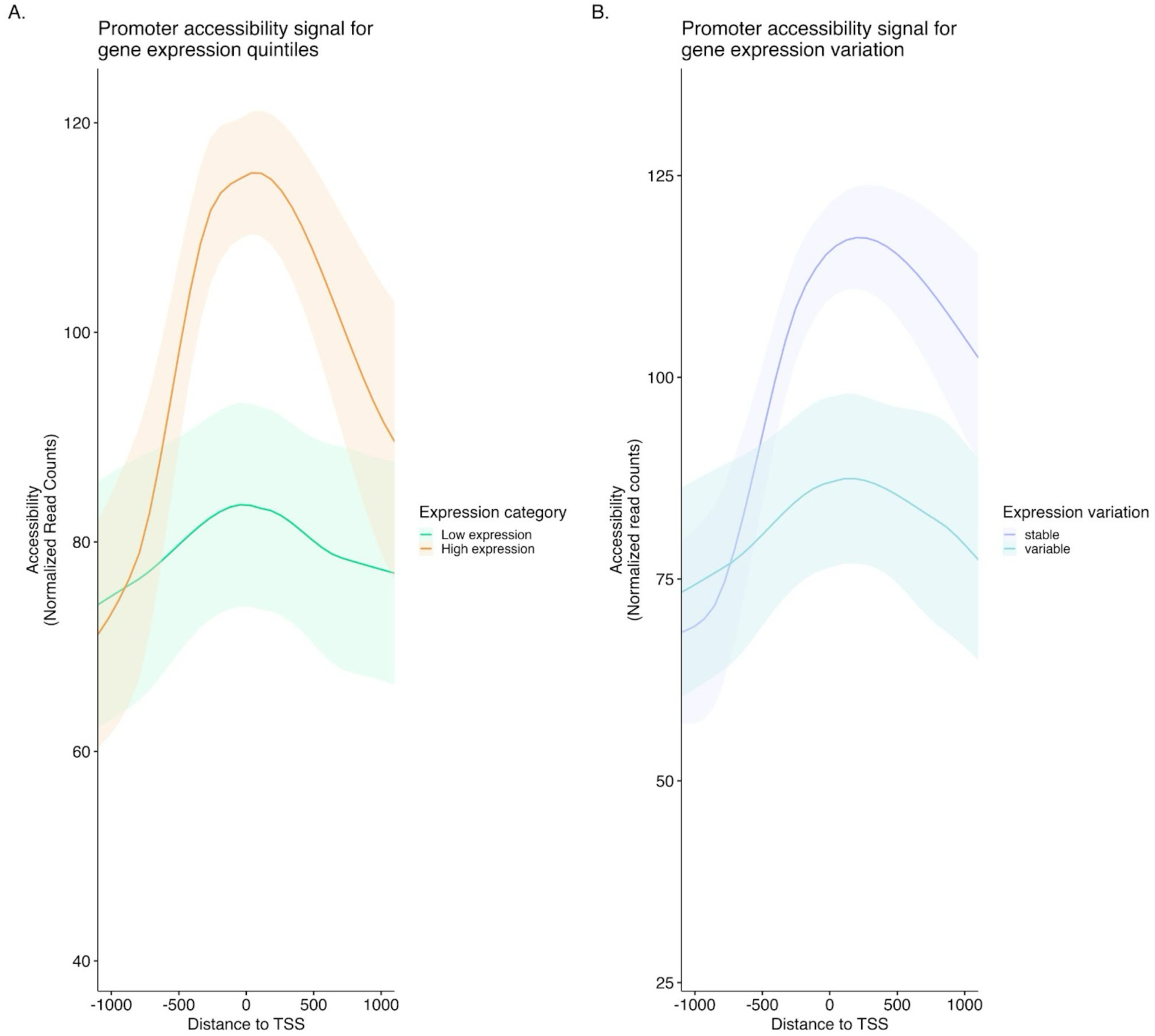
**A. Relationship between promoter accessibility and gene expression.** Genes in the 5th quintile of overall mean expression exhibit the highest chromatin accessibility around the transcription start site (TSS) compared to the genes in lower expression quintiles. **B. Relationship between promoter accessibility and gene expression variation.** Genes with stable expression exhibit higher chromatin accessibility around the TSS compared to genes with variable expression.

Among the 2,227 heat response genes, 201 had promoter-region chromatin accessibility. A Fisher’s exact test comparing the frequency of accessible promoters in heat response genes versus non–heat response genes revealed no significant association (p = 0.763). This lack of enrichment is consistent with our observation that genes exhibiting high expression variability tend to have less accessible promoters (Figure 4). One possible explanation is that many heat response genes are transcriptionally silent under ambient conditions, with promoters that are inaccessible but epigenetically poised for rapid activation during heat stress. Supporting this, we found a weak but significant positive correlation between the mean expression level of heat response genes and promoter accessibility (Figure 5A; Spearman’s ρ = 0.11, p < 0.05). However, no relationship was observed between promoter accessibility and the magnitude of expression change upon heat stress (Figure 5B; Spearman’s p > 0.1), suggesting that baseline chromatin accessibility at promoters does not predict the strength of transcriptional response.

**Figure 5.**
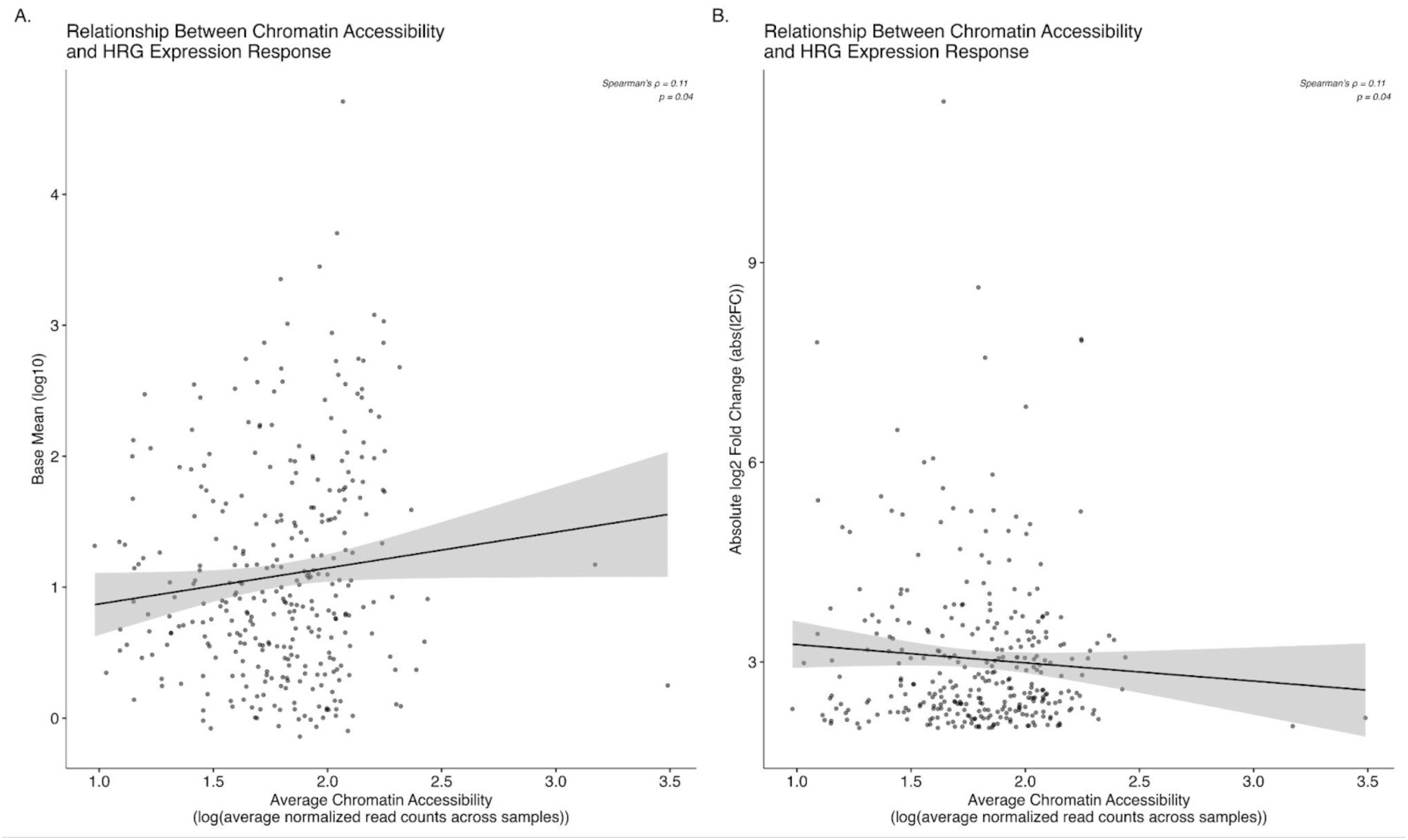
**A. Correlation between mean gene expression of heat response genes and promoter accessibility.** Slight positive relationship between mean expression and baseline promoter accessibility. **B. Correlation between log_2_ fold change of heat response genes and promoter accessibility.** No relationship observed between log2 fold change following heat stress and the baseline promoter accessibility.

## Discussion

Ecological epigenetics aims to understand how non-genetic sources of phenotypic variation influence evolutionary trajectories and improve predictions of species responses to changing climates (Lamka et al. 2022). Previous ecological epigenomic studies have primarily focused on the role of DNA methylation in regulating gene expression for ecologically relevant traits. This focus is driven by the well-established link between DNA methylation and gene expression, the environmentally induced fluctuations in DNA methylation, and the relative ease of generating such data from natural populations (Eirin-Lopez and Putnam 2019; Hofmann 2017; Suzuki and Bird 2008). However, evidence suggests that in various species, fluctuations in DNA methylation play a negligible role in causing short-term changes in gene expression (Abbott et al. 2024; Bewick et al. 2019; Dixon and Matz 2022; Guerrero and Bay 2024). Environmental stressors lead to changes in histone structure in reef building corals, and chromatin accessibility changes are linked with heat exposure in *E. pallida*, suggesting an important role for chromatin organization as a short-term regulator in corals capable of responding to environmental cues (Rodriguez-Casariego et al. 2018; Roquis et al. 2022; Weizman and Levy 2019). In this study, we investigated the relationship between gene expression and chromatin accessibility, contributing to the growing body of knowledge on how species mediate gene expression in response to environmental cues. Our ATAC-seq data revealed a predominance of open chromatin regions in distal intergenic regions, promoters, and introns, highlighting the importance of chromatin accessibility in gene regulation. We also show that genes with accessible chromatin promoters had increased expression and reduced variability. Heat response genes with open promoters on average were upregulated, suggesting that these genes are primed for an immediate heat stress response. Since genes with high expression variability and closed promoters are likely poised to respond to environmental cues, future research should focus on characterizing the mechanisms that enable this potential for gene expression plasticity.

### Phenotypic and Transcriptional Responses to Heat Stress

Heat stress impacted both symbiont and coral homeostasis, yet thermal history did not significantly affect the physiological or gene expression responses to heat stress. Previous studies have found that *Acropora* corals exhibit diminished sensitivity to higher temperatures following exposure to warmer conditions, a response that is influenced by the duration of exposure and the time elapsed since that exposure (Bay and Palumbi 2015; Bellantuono et al. 2012; Middlebrook et al. 2008). For example, a previous study on *A. nana* found that an elevated temperature of just 2°C for 7 days led to increased tolerance to acute heat stress (Bay and Palumbi 2015). Under similar acclimation conditions, we saw no difference in thermal tolerance based on thermal history in *A. millepora*. One possible explanation is that the acclimation temperature was not warm enough to induce an acclimation response. Alternatively, the corals used in our experiment, which were raised in thermally stable aquaria, have lost the plastic capacity of wild-raised corals and thus do not exhibit the strong acclimation response seen in other studies. The functional enrichment of GO terms associated with heat response genes aligns with previous findings on the essential cellular processes involved in responding to heat stress (Bellantuono et al. 2012; Granados-Cifuentes et al. 2013; Louis et al. 2017).

### Open chromatin regions in the *A. millepora* genome

We used ATAC-seq to map open chromatin regions in the *A. millepora* genome (Buenrostro et al. 2013; Buenrostro et al. 2015). Chromatin accessibility is an epigenetic layer that interacts with other regulatory mechanisms. Its role in mediating gene expression in response to environmental cues is increasingly being studied in ecologically relevant species (Klemm et al. 2019; Lamka et al. 2022; Turner 2009). In our study, open distal regions constituted the majority of the identified peaks, which is consistent with the higher proportion of distal intergenic regions typically reported in ATAC-seq experiments (Yan et al. 2020). This contrasts with results in the cnidarian model *Exaiptasia pallida*, which demonstrate lower proportion of intergenic distal accessible regions, suggesting species-specific differences (Weizman and Levy 2019). Open distal chromatin regions are increasingly recognized for their role in gene expression regulation and are hypothesized to be evolutionarily significant (Pliner et al. 2018; Roscito et al. 2018; Weizman and Levy 2019; Yan et al. 2020; Horvath et al. 2021). For example, open intergenic chromatin regions in the plant species, *Capsella grandiflora*, serve as a reservoir for mutations with regulatory functions, often having minimal fitness effects, thereby allowing them to accumulate over time (Horvath et al. 2021). Further exploration of genetic variation within open distal regions in *A. millepora* and other ecologically relevant species could shed light on the evolutionary mechanisms that shape gene regulation variation.

Functional enrichment signifies that genes with open chromatin promoters in ambient conditions are involved in regulating immune responses, viral processes, and gene silencing mechanisms, indicating their role in controlling essential cellular operations. Open promoters facilitate binding of transcriptional activators, enhancing gene expression (Cairns 2009). Enrichment of genes with open promoters, such as those involved in cellular metabolic processes and RNA nuclease activity, suggest that accessibility of these promoters is important to maintaining baseline cellular functions. Conversely, the enrichment of terms related to immune and external stimuli responses suggests that promoters are poised to rapidly respond to environmental cues (Puri et al. 2015). Specifically, the presence of positive and negative regulation of immune system processes at baseline suggests a finely tuned system that maintains immune stability while being prepared for activation. Similarities in functional enrichment of genes with open promoters between symbiotic *E. pallida* and *A. millepora* such as regulation of gene expression and responses to stress and stimuli suggest that some patterns of chromatin accessibility may be conserved across cnidarians (Weizman and Levy 2019).

Enriched regulatory processes observed in ambient conditions may reveal novel gene expression mechanisms, offering insights into important pathways that could explain adaptive processes in coral, which have yet to be fully explored (Eirin-Lopez and Putnam 2019). For instance, the enrichment for gene silencing by regulation of non-coding RNA (ncRNA) and microRNA (miRNA) supports the growing recognition that regulatory RNAs are critical for stress responses in corals (Huang et al. 2019; Liew et al. 2014). Studies across taxa demonstrate that ncRNAs play specific roles in responding to stressors such as temperature and may contribute to gene expression changes linked to phenotypic plasticity (Heo and Sung 2011; Kindgren et al. 2018; Eirin-Lopez and Putnam 2019; Jha et al. 2023). Further investigation into these regulatory mechanisms in ecologically relevant species will be critical for understanding how these organisms adapt to changing environments and represents an exciting avenue for future research.

### Relationship Between Open Chromatin Promoters and Gene Expression

In our study, genes with the highest expression and lowest variability exhibited high chromatin accessibility around the TSS. This relationship, also observed in marbled crayfish, is closely associated with gene-body DNA methylation (Gatzmann et al. 2018). Genes with high expression and promoter accessibility were depleted of gene-body methylation, while heavy methylation was linked to reduced promoter accessibility in stably expressed genes, suggesting a role in maintaining transcriptional consistency. Although we did not assess DNA methylation here, relationships observed in other invertebrates, including Acropora, could inform hypotheses about its interaction with chromatin accessibility in regulating coral gene expression. In invertebrates, housekeeping genes—stably expressed to maintain homeostasis—are typically more densely methylated than environmentally responsive genes (Dixon and Matz 2022; Gatzmann et al. 2018). Moreover, gene expression variability, reflecting noise and environmental response, tends to decrease as gene-body methylation increases (Dixon and Matz 2022; Gatzmann et al. 2018; Guerrero and Bay 2024). These findings support the hypothesis that chromatin accessibility and DNA methylation synergistically mediate gene expression homeostasis (Gatzmann et al. 2018; Li et al. 2018).

It is compelling to speculate on the histone modifications that may influence the chromatin state of housekeeping and environmentally responsive genes. Our finding that highly expressed genes exhibit the greatest promoter accessibility is consistent with mouse embryonic stem cells (Clark et al. 2018). In mouse fibroblasts, H4K20me1 correlates with chromatin accessibility and expression, particularly in housekeeping genes (Shoaib et al. 2021). Histone H4 is highly conserved between cnidarians and mice, suggesting these mechanisms may be evolutionarily conserved (Roquis et al. 2022). Yet epigenetic layers, such as DNA methylation, differ: vertebrate genomes are densely methylated except in regulatory regions (Klughammer et al. 2023; Suzuki and Bird 2008), whereas invertebrate genomes are sparsely methylated, with gene body–biased methylation positively associated with expression (Dixon and Matz 2022; Roberts and Gavery 2012). Thus, while some histone modifications may be conserved, their interactions with other epigenetic factors vary across taxa.

Mammalian oocytes, preimplantation embryos, and placenta display methylation landscapes more similar to invertebrates (Demond and Kelsey 2020; Smallwood et al. 2011), providing insight into conserved histone regulation. In these sparsely methylated tissues, poised chromatin marks essential for development include H3K4me3, H3K27ac, and H2A.Z modifications (Dijkwel and Tremethick 2022; Sotomayor-Lugo et al. 2024). For instance, when the H3K4me3 footprint around the TSS is wider than 5kb, this histone mark is associated with activated transcription and maintains stable DNA methylation percentages (Liu et al. 2016; Sotomayor-Lugo et al. 2024). Co-occurrence of H3K4me3 and H3K27ac marks transient, stage-specific expression (Sotomayor-Lugo et al. 2024; Wu et al. 2023). Chromatin accessibility in H3K4me3-marked regions depends on footprint breadth and protein associations, yielding context-dependent structure (Beacon et al. 2021). These findings suggest conserved roles for histone modifications in linking gene expression, DNA methylation, and chromatin accessibility, though their interplay remains complex and requires further study in corals and other invertebrates (Beck et al. 2012; Corvalan and Coller 2021).

Accessible chromatin promoters established under ambient conditions appear to play a small but significant role in mediating gene expression plasticity required to respond to heat stress. Promoters can be poised to repress expression under ambient conditions while simultaneously remaining primed to rapidly respond to various cellular stimuli (Puri et al. 2015). In our study, we identified a set of heat response genes with promoters poised to respond to heat stress. These genes were primarily upregulated compared to heat response genes with closed promoters in ambient conditions. A recent study by Himanen et al. (2022) demonstrated that heat shock transcription factors (HSFs) can reprogram transcription by either directly or indirectly facilitating the release of paused RNA polymerase II in open chromatin near the TSS. HSFs are also known to play a crucial role in coral thermal-tolerance (Cleves et al. 2020; Hayes and King 1995; Ishii et al. 2019; Meyer et al. 2011). In our study, functional enrichment analyses of the poised heat response genes suggest that they are involved in high-level regulatory processes, including gene expression and protein modifications. The essential cellular cascades regulated by heat response genes with poised promoters are linked to immune responses, stress responses, apoptosis, and post-translational modifications. These findings offer insight into how transcription initiation may occur in genes with poised promoters during heat stress in ecologically relevant species.

