## Supplement table 1 for "Chromatin accessibility and heat stress gene expression in the reef-building coral, *Acropora millepora*"

**Gene Ontology (GO) terms of heat response genes**

Table shows the results from a GO enrichment analysis of heat response genes using the R package, topGO v. 2.50.0. We compared the list of genes with significant differential expression against a background of all *A. millepora* annotated transcripts. Enriched GO terms were identified with the classic Fisher's test with a p-value < 0.01 and at least 10 genes within each category. GO terms are contained within ontology types: Biological Processes (BP), Cellular Components (CC), and Molecular Functions (MF).

| Type | GO ID | Term | p-value |
|---|---|---|---|
| MF | GO:0016616 | oxidoreductase activity, acting on the CH-OH group of donors, NAD or NADP as acceptor | 0.00011 |
| MF | GO:0005342 | organic acid transmembrane transporter activity | 0.00011 |
| MF | GO:0046943 | carboxylic acid transmembrane transporter activity | 0.00011 |
| MF | GO:0030545 | signaling receptor regulator activity | 0.00017 |
| MF | GO:0005343 | organic acid:sodium symporter activity | 0.00018 |
| MF | GO:0016846 | carbon-sulfur lyase activity | 0.00018 |
| MF | GO:0048018 | receptor ligand activity | 0.00032 |
| MF | GO:0015294 | solute:monoatomic cation symporter activity | 0.00037 |
| MF | GO:0030546 | signaling receptor activator activity | 0.00045 |
| MF | GO:0000977 | RNA polymerase II transcription regulatory region sequence-specific DNA binding | 0.00047 |
| MF | GO:0015370 | solute:sodium symporter activity | 0.00048 |
| MF | GO:0008514 | organic anion transmembrane transporter activity | 0.00072 |
| MF | GO:0015171 | amino acid transmembrane transporter activity | 0.00076 |
| MF | GO:0015293 | symporter activity | 0.00084 |
| MF | GO:0046872 | metal ion binding | 0.0011 |
| MF | GO:0000976 | transcription cis-regulatory region binding | 0.00114 |
| MF | GO:0001067 | transcription regulatory region nucleic acid binding | 0.00114 |
| MF | GO:0008233 | peptidase activity | 0.0012 |
| MF | GO:0015179 | L-amino acid transmembrane transporter activity | 0.00131 |
| MF | GO:0005201 | extracellular matrix structural constituent | 0.00136 |
| MF | GO:0008238 | exopeptidase activity | 0.00138 |
| MF | GO:0016836 | hydro-lyase activity | 0.0015 |
| MF | GO:0051082 | unfolded protein binding | 0.00159 |
| MF | GO:0016209 | antioxidant activity | 0.00188 |
| MF | GO:0043169 | cation binding | 0.00204 |
| MF | GO:0008083 | growth factor activity | 0.00224 |
| MF | GO:0008235 | metalloexopeptidase activity | 0.00304 |
| MF | GO:0005506 | iron ion binding | 0.00331 |
| MF | GO:0016705 | oxidoreductase activity, acting on paired donors, with incorporation or reduction of molecular oxyge... | 0.00436 |
| MF | GO:1990837 | sequence-specific double-stranded DNA binding | 0.00491 |
| MF | GO:0043565 | sequence-specific DNA binding | 0.00531 |
| MF | GO:0015291 | secondary active transmembrane transporter activity | 0.00533 |

| | | | |
|---|---|---|---|
| MF | GO:0038024 | cargo receptor activity | 0.00548 |
| MF | GO:0000978 | RNA polymerase II cis-regulatory region sequence-specific DNA binding | 0.00623 |
| MF | GO:0000987 | cis-regulatory region sequence-specific DNA binding | 0.00701 |
| MF | GO:0004180 | carboxypeptidase activity | 0.00945 |
| BP | GO:0012501 | programmed cell death | 0.0001 |
| BP | GO:0007166 | cell surface receptor signaling pathway | 0.0001 |
| BP | GO:0008202 | steroid metabolic process | 0.00011 |
| BP | GO:0046638 | positive regulation of alpha-beta T cell differentiation | 0.00011 |
| BP | GO:0001525 | angiogenesis | 0.00011 |
| BP | GO:0002521 | leukocyte differentiation | 0.00011 |
| BP | GO:0002819 | regulation of adaptive immune response | 0.00011 |
| BP | GO:0002376 | immune system process | 0.00012 |
| BP | GO:0009893 | positive regulation of metabolic process | 0.00012 |
| BP | GO:0051094 | positive regulation of developmental process | 0.00012 |
| BP | GO:0045860 | positive regulation of protein kinase activity | 0.00013 |
| BP | GO:0042325 | regulation of phosphorylation | 0.00014 |
| BP | GO:0046637 | regulation of alpha-beta T cell differentiation | 0.00014 |
| BP | GO:0010604 | positive regulation of macromolecule metabolic process | 0.00014 |
| BP | GO:0015849 | organic acid transport | 0.00015 |
| BP | GO:0046942 | carboxylic acid transport | 0.00015 |
| BP | GO:0046634 | regulation of alpha-beta T cell activation | 0.00015 |
| BP | GO:0033674 | positive regulation of kinase activity | 0.00016 |
| BP | GO:0045621 | positive regulation of lymphocyte differentiation | 0.00016 |
| BP | GO:0006749 | glutathione metabolic process | 0.00016 |
| BP | GO:0002292 | T cell differentiation involved in immune response | 0.00017 |
| BP | GO:0002822 | regulation of adaptive immune response based on somatic recombination of immune receptors built from... | 0.00019 |
| BP | GO:0046631 | alpha-beta T cell activation | 0.0002 |
| BP | GO:0045596 | negative regulation of cell differentiation | 0.0002 |
| BP | GO:0002252 | immune effector process | 0.0002 |
| BP | GO:0002449 | lymphocyte mediated immunity | 0.00021 |
| BP | GO:1903131 | mononuclear cell differentiation | 0.00022 |
| BP | GO:0048522 | positive regulation of cellular process | 0.00026 |
| BP | GO:0007159 | leukocyte cell-cell adhesion | 0.00028 |
| BP | GO:0050863 | regulation of T cell activation | 0.00029 |
| BP | GO:1903037 | regulation of leukocyte cell-cell adhesion | 0.0003 |
| BP | GO:1903706 | regulation of hemopoiesis | 0.00031 |
| BP | GO:0019220 | regulation of phosphate metabolic process | 0.00033 |
| BP | GO:0071900 | regulation of protein serine/threonine kinase activity | 0.00033 |
| BP | GO:0051174 | regulation of phosphorus metabolic process | 0.00035 |
| BP | GO:0002711 | positive regulation of T cell mediated immunity | 0.00037 |
| BP | GO:0045595 | regulation of cell differentiation | 0.00037 |
| BP | GO:1901700 | response to oxygen-containing compound | 0.00038 |
| BP | GO:0042221 | response to chemical | 0.00038 |
| BP | GO:0002443 | leukocyte mediated immunity | 0.00038 |
| BP | GO:0050866 | negative regulation of cell activation | 0.0004 |
| BP | GO:0007154 | cell communication | 0.0004 |
| BP | GO:0006468 | protein phosphorylation | 0.0004 |
| BP | GO:0060429 | epithelium development | 0.00041 |
| BP | GO:0032502 | developmental process | 0.00041 |
| BP | GO:0007167 | enzyme-linked receptor protein signaling pathway | 0.00042 |
| BP | GO:0050865 | regulation of cell activation | 0.00043 |
| BP | GO:0071363 | cellular response to growth factor stimulus | 0.00043 |
| BP | GO:0030856 | regulation of epithelial cell differentiation | 0.00044 |
| BP | GO:0048856 | anatomical structure development | 0.00044 |
| BP | GO:0001934 | positive regulation of protein phosphorylation | 0.00048 |
| BP | GO:0002286 | T cell activation involved in immune response | 0.00049 |
| BP | GO:0051338 | regulation of transferase activity | 0.00051 |
| BP | GO:0048646 | anatomical structure formation involved in morphogenesis | 0.00052 |
| BP | GO:0051249 | regulation of lymphocyte activation | 0.00053 |
| BP | GO:0009628 | response to abiotic stimulus | 0.00056 |
| BP | GO:0031325 | positive regulation of cellular metabolic process | 0.00056 |

| | | | |
|---|---|---|---|
| BP | GO:0007169 | transmembrane receptor protein tyrosine kinase signaling pathway | 0.00057 |
| BP | GO:0043405 | regulation of MAP kinase activity | 0.00057 |
| BP | GO:0015711 | organic anion transport | 0.00061 |
| BP | GO:0006955 | immune response | 0.00064 |
| BP | GO:0002250 | adaptive immune response | 0.00065 |
| BP | GO:0046323 | glucose import | 0.00067 |
| BP | GO:0002768 | immune response-regulating cell surface receptor signaling pathway | 0.00068 |
| BP | GO:0060021 | roof of mouth development | 0.00069 |
| BP | GO:0009636 | response to toxic substance | 0.00069 |
| BP | GO:0042981 | regulation of apoptotic process | 0.00073 |
| BP | GO:0042327 | positive regulation of phosphorylation | 0.00077 |
| BP | GO:0070848 | response to growth factor | 0.0008 |
| BP | GO:0001503 | ossification | 0.00083 |
| BP | GO:0042446 | hormone biosynthetic process | 0.00084 |
| BP | GO:0007165 | signal transduction | 0.00086 |
| BP | GO:0072006 | nephron development | 0.00096 |
| BP | GO:0002709 | regulation of T cell mediated immunity | 0.00097 |
| BP | GO:0016310 | phosphorylation | 0.00102 |
| BP | GO:0010562 | positive regulation of phosphorus metabolic process | 0.00106 |
| BP | GO:0045937 | positive regulation of phosphate metabolic process | 0.00106 |
| BP | GO:0043067 | regulation of programmed cell death | 0.00107 |
| BP | GO:0061448 | connective tissue development | 0.00108 |
| BP | GO:0051251 | positive regulation of lymphocyte activation | 0.00109 |
| BP | GO:0002366 | leukocyte activation involved in immune response | 0.0011 |
| BP | GO:0022409 | positive regulation of cell-cell adhesion | 0.00112 |
| BP | GO:0032743 | positive regulation of interleukin-2 production | 0.00116 |
| BP | GO:0055085 | transmembrane transport | 0.0012 |
| BP | GO:0042743 | hydrogen peroxide metabolic process | 0.00121 |
| BP | GO:0071466 | cellular response to xenobiotic stimulus | 0.00121 |
| BP | GO:1902106 | negative regulation of leukocyte differentiation | 0.00121 |
| BP | GO:0002263 | cell activation involved in immune response | 0.00123 |
| BP | GO:0002429 | immune response-activating cell surface receptor signaling pathway | 0.00124 |
| BP | GO:0033993 | response to lipid | 0.00126 |
| BP | GO:0090287 | regulation of cellular response to growth factor stimulus | 0.00127 |
| BP | GO:0002694 | regulation of leukocyte activation | 0.00131 |
| BP | GO:0032147 | activation of protein kinase activity | 0.00131 |
| BP | GO:0007155 | cell adhesion | 0.00134 |
| BP | GO:0051347 | positive regulation of transferase activity | 0.00136 |
| BP | GO:0031328 | positive regulation of cellular biosynthetic process | 0.00147 |
| BP | GO:1902074 | response to salt | 0.00148 |
| BP | GO:0023052 | signaling | 0.00149 |
| BP | GO:0002456 | T cell mediated immunity | 0.00149 |
| BP | GO:0002824 | positive regulation of adaptive immune response based on somatic recombination of immune receptors b... | 0.00158 |
| BP | GO:1903707 | negative regulation of hemopoiesis | 0.0016 |
| BP | GO:0015698 | inorganic anion transport | 0.00162 |
| BP | GO:0045785 | positive regulation of cell adhesion | 0.00163 |
| BP | GO:0032663 | regulation of interleukin-2 production | 0.00169 |
| BP | GO:0000122 | negative regulation of transcription by RNA polymerase II | 0.00174 |
| BP | GO:0045444 | fat cell differentiation | 0.00176 |
| BP | GO:0071902 | positive regulation of protein serine/threonine kinase activity | 0.00204 |
| BP | GO:0044344 | cellular response to fibroblast growth factor stimulus | 0.00204 |
| BP | GO:0072593 | reactive oxygen species metabolic process | 0.00208 |
| BP | GO:0032623 | interleukin-2 production | 0.0021 |
| BP | GO:0009891 | positive regulation of biosynthetic process | 0.00212 |
| BP | GO:0002708 | positive regulation of lymphocyte mediated immunity | 0.00229 |
| BP | GO:0002821 | positive regulation of adaptive immune response | 0.00229 |
| BP | GO:0031214 | biomineral tissue development | 0.0023 |
| BP | GO:0002699 | positive regulation of immune effector process | 0.00234 |
| BP | GO:0035556 | intracellular signal transduction | 0.00238 |
| BP | GO:0002040 | sprouting angiogenesis | 0.00257 |
| BP | GO:0003073 | regulation of systemic arterial blood pressure | 0.00265 |

| | | | |
|---|---|---|---|
| BP | GO:0009605 | response to external stimulus | 0.00271 |
| BP | GO:0002705 | positive regulation of leukocyte mediated immunity | 0.00274 |
| BP | GO:0030183 | B cell differentiation | 0.00274 |
| BP | GO:0030154 | cell differentiation | 0.00286 |
| BP | GO:0010557 | positive regulation of macromolecule biosynthetic process | 0.00292 |
| BP | GO:0048869 | cellular developmental process | 0.00293 |
| BP | GO:0070887 | cellular response to chemical stimulus | 0.00294 |
| BP | GO:0002220 | innate immune response activating cell surface receptor signaling pathway | 0.00296 |
| BP | GO:0051250 | negative regulation of lymphocyte activation | 0.00296 |
| BP | GO:0006950 | response to stress | 0.003 |
| BP | GO:0032496 | response to lipopolysaccharide | 0.00303 |
| BP | GO:0031401 | positive regulation of protein modification process | 0.00303 |
| BP | GO:0006865 | amino acid transport | 0.00308 |
| BP | GO:0030097 | hemopoiesis | 0.00309 |
| BP | GO:0002695 | negative regulation of leukocyte activation | 0.00316 |
| BP | GO:0001101 | response to acid chemical | 0.0032 |
| BP | GO:0016125 | sterol metabolic process | 0.00327 |
| BP | GO:0045597 | positive regulation of cell differentiation | 0.00332 |
| BP | GO:0010035 | response to inorganic substance | 0.0034 |
| BP | GO:0043066 | negative regulation of apoptotic process | 0.00341 |
| BP | GO:0045765 | regulation of angiogenesis | 0.00344 |
| BP | GO:0002682 | regulation of immune system process | 0.00348 |
| BP | GO:0070661 | leukocyte proliferation | 0.00351 |
| BP | GO:0045944 | positive regulation of transcription by RNA polymerase II | 0.00356 |
| BP | GO:0006357 | regulation of transcription by RNA polymerase II | 0.00374 |
| BP | GO:0001657 | ureteric bud development | 0.00375 |
| BP | GO:0009064 | glutamine family amino acid metabolic process | 0.00375 |
| BP | GO:0008543 | fibroblast growth factor receptor signaling pathway | 0.00377 |
| BP | GO:0030282 | bone mineralization | 0.00382 |
| BP | GO:0019221 | cytokine-mediated signaling pathway | 0.00382 |
| BP | GO:0071774 | response to fibroblast growth factor | 0.00388 |
| BP | GO:0002237 | response to molecule of bacterial origin | 0.00388 |
| BP | GO:0016477 | cell migration | 0.00393 |
| BP | GO:0048583 | regulation of response to stimulus | 0.00396 |
| BP | GO:0034764 | positive regulation of transmembrane transport | 0.004 |
| BP | GO:0051716 | cellular response to stimulus | 0.00405 |
| BP | GO:0097190 | apoptotic signaling pathway | 0.00406 |
| BP | GO:0072080 | nephron tubule development | 0.00419 |
| BP | GO:0009410 | response to xenobiotic stimulus | 0.00419 |
| BP | GO:0060324 | face development | 0.00419 |
| BP | GO:0072073 | kidney epithelium development | 0.00419 |
| BP | GO:0002696 | positive regulation of leukocyte activation | 0.0042 |
| BP | GO:0042476 | odontogenesis | 0.00437 |
| BP | GO:0001658 | branching involved in ureteric bud morphogenesis | 0.00448 |
| BP | GO:0006952 | defense response | 0.0045 |
| BP | GO:0051402 | neuron apoptotic process | 0.0045 |
| BP | GO:0035264 | multicellular organism growth | 0.00456 |
| BP | GO:0015807 | L-amino acid transport | 0.0046 |
| BP | GO:0048880 | sensory system development | 0.00464 |
| BP | GO:0031399 | regulation of protein modification process | 0.00469 |
| BP | GO:1901342 | regulation of vasculature development | 0.0047 |
| BP | GO:0043069 | negative regulation of programmed cell death | 0.00475 |
| BP | GO:0051246 | regulation of protein metabolic process | 0.00484 |
| BP | GO:1902652 | secondary alcohol metabolic process | 0.00487 |
| BP | GO:0001935 | endothelial cell proliferation | 0.00522 |
| BP | GO:0043406 | positive regulation of MAP kinase activity | 0.00529 |
| BP | GO:0050867 | positive regulation of cell activation | 0.00533 |
| BP | GO:0051781 | positive regulation of cell division | 0.00543 |
| BP | GO:0001823 | mesonephros development | 0.00553 |
| BP | GO:0072163 | mesonephric epithelium development | 0.00553 |
| BP | GO:0072164 | mesonephric tubule development | 0.00553 |
| BP | GO:0009725 | response to hormone | 0.00555 |

| | | | |
|---|---|---|---|
| BP | GO:0048585 | negative regulation of response to stimulus | 0.00601 |
| BP | GO:0008217 | regulation of blood pressure | 0.00616 |
| BP | GO:0008203 | cholesterol metabolic process | 0.00636 |
| BP | GO:0009314 | response to radiation | 0.00637 |
| BP | GO:0046903 | secretion | 0.00641 |
| BP | GO:0009617 | response to bacterium | 0.00658 |
| BP | GO:0006694 | steroid biosynthetic process | 0.00676 |
| BP | GO:0072009 | nephron epithelium development | 0.00676 |
| BP | GO:0042127 | regulation of cell population proliferation | 0.00679 |
| BP | GO:0002697 | regulation of immune effector process | 0.00696 |
| BP | GO:0048705 | skeletal system morphogenesis | 0.00696 |
| BP | GO:0005996 | monosaccharide metabolic process | 0.00696 |
| BP | GO:0040011 | locomotion | 0.00704 |
| BP | GO:1901701 | cellular response to oxygen-containing compound | 0.00705 |
| BP | GO:0045862 | positive regulation of proteolysis | 0.00725 |
| BP | GO:0010033 | response to organic substance | 0.00726 |
| BP | GO:0006665 | sphingolipid metabolic process | 0.00726 |
| BP | GO:0034097 | response to cytokine | 0.00728 |
| BP | GO:0001654 | eye development | 0.00741 |
| BP | GO:0150063 | visual system development | 0.00741 |
| BP | GO:0045321 | leukocyte activation | 0.00764 |
| BP | GO:0050851 | antigen receptor-mediated signaling pathway | 0.00777 |
| BP | GO:0048468 | cell development | 0.00781 |
| BP | GO:0060675 | ureteric bud morphogenesis | 0.00783 |
| BP | GO:0072171 | mesonephric tubule morphogenesis | 0.00783 |
| BP | GO:0050776 | regulation of immune response | 0.00799 |
| BP | GO:0045893 | positive regulation of DNA-templated transcription | 0.00815 |
| BP | GO:0051254 | positive regulation of RNA metabolic process | 0.00823 |
| BP | GO:0002706 | regulation of lymphocyte mediated immunity | 0.00825 |
| BP | GO:0001656 | metanephros development | 0.00826 |
| BP | GO:0022407 | regulation of cell-cell adhesion | 0.00835 |
| BP | GO:0050790 | regulation of catalytic activity | 0.00836 |
| BP | GO:0006629 | lipid metabolic process | 0.00846 |
| BP | GO:1902680 | positive regulation of RNA biosynthetic process | 0.00849 |
| BP | GO:0048562 | embryonic organ morphogenesis | 0.00867 |
| BP | GO:0002684 | positive regulation of immune system process | 0.0088 |
| BP | GO:0009790 | embryo development | 0.00886 |
| BP | GO:0071356 | cellular response to tumor necrosis factor | 0.00906 |
| BP | GO:0048598 | embryonic morphogenesis | 0.00933 |
| BP | GO:0044281 | small molecule metabolic process | 0.00938 |
| BP | GO:0046777 | protein autophosphorylation | 0.00984 |
| BP | GO:0009968 | negative regulation of signal transduction | 0.00989 |
| BP | GO:0001817 | regulation of cytokine production | 0.00997 |
| CC | GO:0045177 | apical part of cell | 0.00011 |
| CC | GO:0005615 | extracellular space | 0.00023 |
| CC | GO:0016324 | apical plasma membrane | 0.0004 |
| CC | GO:0071944 | cell periphery | 0.00098 |
| CC | GO:0005886 | plasma membrane | 0.00246 |
| CC | GO:0016020 | membrane | 0.004 |
| CC | GO:0005581 | collagen trimer | 0.00401 |
| CC | GO:0005764 | lysosome | 0.00538 |
| CC | GO:0005796 | Golgi lumen | 0.00846 |
