## Supplement figure 1 for "Chromatin accessibility and heat stress gene expression in the reef-building coral, *Acropora millepora*"

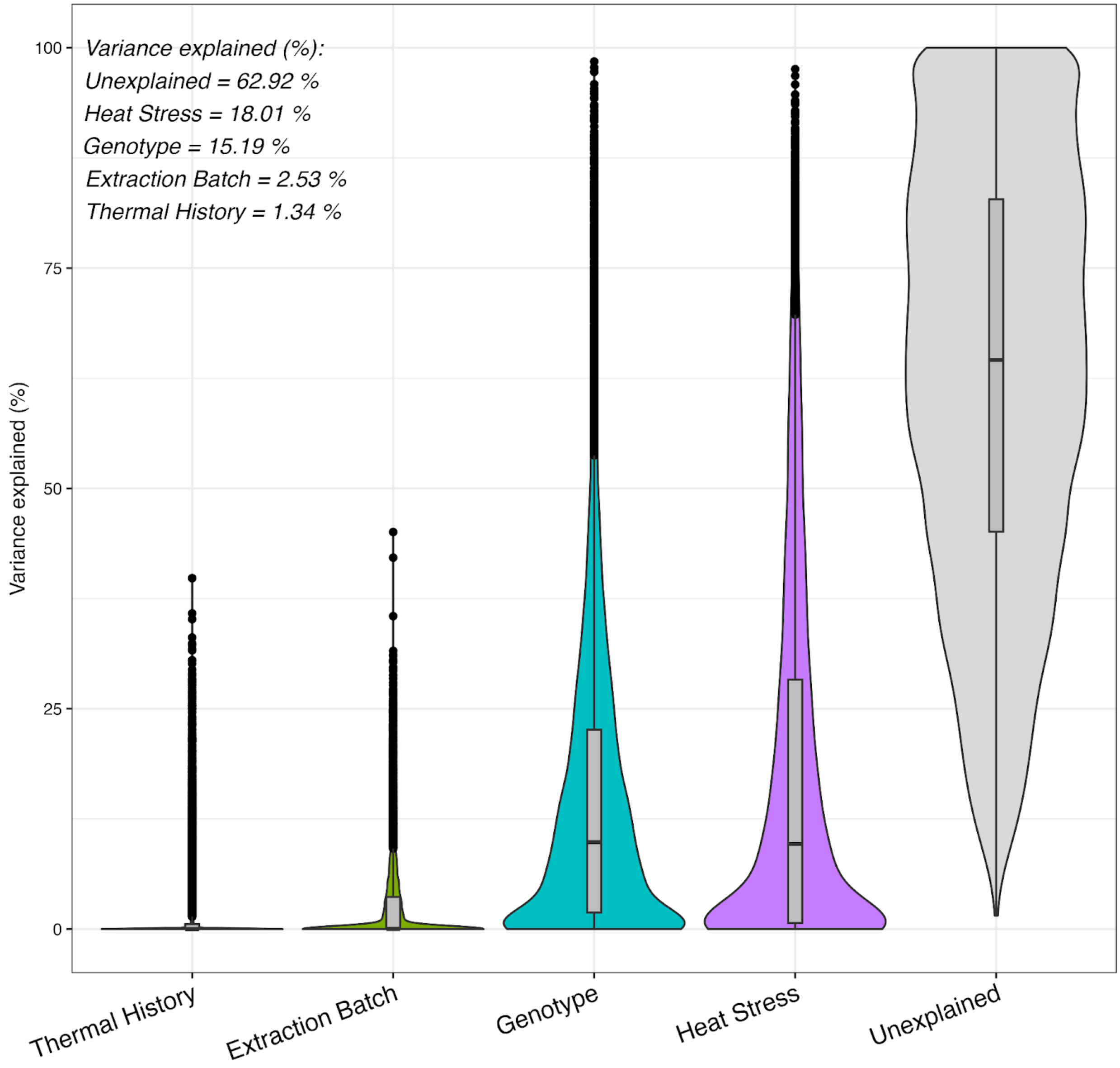


**Variance partitioning of gene expression across covariates** Variation in thermal history among coral fragments did not result in measurable differences in gene expression following the heat-stress assay. This indicates that our incubation strategy did not elicit the intended acclimatory effect.
